# The unconventional kinetoplastid kinetochore protein KKT4 tracks with dynamic microtubule tips

**DOI:** 10.1101/216812

**Authors:** Aida Llauró, Hanako Hayashi, Megan E. Bailey, Alex Wilson, Patryk Ludzia, Charles L. Asbury, Bungo Akiyoshi

## Abstract

Kinetochores are multiprotein machines that drive chromosome segregation in all eukaryotes by maintaining persistent, load-bearing linkages between the chromosomes and the tips of dynamic spindle microtubules. Kinetochores in commonly studied eukaryotes are assembled from widely conserved components like the Ndc80 complex that directly binds microtubules. However, in evolutionarily-divergent kinetoplastid species such as *Trypanosoma brucei*, which causes sleeping sickness, the kinetochores assemble from a unique set of proteins lacking homology to any known microtubule-binding domains. Here we show that a kinetochore protein from *T. brucei* called KKT4 binds directly to microtubules, diffuses along the microtubule lattice, and tracks with disassembling microtubule tips. The protein localizes both to kinetochores and to spindle microtubules in vivo, and its depletion causes defects in chromosome segregation. We define a minimal microtubule-binding domain within KKT4 and identify several charged residues important for its microtubule-binding activity. Laser trapping experiments show that KKT4 can maintain load-bearing attachments to both growing and shortening microtubule tips. Thus, despite its lack of similarity to other known microtubule-binding proteins, KKT4 has key functions required for harnessing microtubule dynamics to drive chromosome segregation. We propose that it represents a primary element of the kinetochore-microtubule interface in kinetoplastids.

## Results and Discussions

Chromosome segregation in eukaryotes depends on the interaction between chromosomes and dynamic spindle microtubules (McIntosh, 2016). The interaction is mediated by the macromolecular kinetochore complex that assembles onto the centromeric region of each chromosome (Cheeseman, 2014; Musacchio and Desai, 2017). Spindle microtubules are dynamic polymers that grow and shrink by addition and loss of tubulin subunits from their tips (Desai and Mitchison, 1997). Accurate chromosome segregation requires that kinetochores maintain persistent, load-bearing attachments to dynamic microtubule tips, even as the tips assemble and disassemble under their grip (Joglekar et al., 2010; McIntosh, 2017). The kinetochore consists of more than 30 structural proteins even in a simple budding yeast kinetochore (Biggins, 2013). Among these components, CENP-A is a centromere-specific histone H3 variant that forms a specialized chromatin environment at the centromere, while the Ndc80 complex directly binds microtubules to mediate the coupling of kinetochores to dynamic microtubule tips (Musacchio and Desai, 2017). The Ndc80 complex consists of Ndc80, Nuf2, Spc24, and Spc25 components (Wigge and Kilmartin, 2001). Microtubule-binding activities reside in the Ndc80 and Nuf2 proteins that have a calponin homology (CH) domain (Cheeseman et al., 2006; Wei et al., 2007; Ciferri et al., 2008). Ensembles of Ndc80 complexes can form load-bearing attachments to dynamic microtubule tips in vitro (McIntosh et al., 2008; Powers et al., 2009). Besides Ndc80, there are several microtubule-binding proteins that localize at kinetochores, including the Dam1 complex and Stu2 protein in yeasts, and the Ska1 complex in metazoa (Cheeseman et al., 2001; He et al., 2001; Tanaka et al., 2005; Hanisch et al., 2006; Hsu and Toda, 2011). These complexes are also capable of forming dynamic, load-bearing attachments in vitro (Asbury et al., 2006; Westermann et al., 2006; Grishchuk et al., 2008; Welburn et al., 2009; Miller et al., 2016). Extensive studies have been carried out to understand the nature of their microtubule-binding properties (Al-Bassam et al., 2006, 2012; Ciferri et al., 2008; Tien et al., 2010; Jeyaprakash et al., 2012; Umbreit et al., 2014; Kim et al., 2017).

Putative homologs of CENP-A and Ndc80 complex components are found in various eukaryotes sequenced thus far (Meraldi et al., 2006; Drinnenberg and Akiyoshi, 2017; van Hooff et al., 2017), suggesting that the majority of eukaryotes use CENP-A and Ndc80 to bind DNA and microtubules, respectively. However, none of these conventional kinetochore proteins has been found in kinetoplastids, a group of evolutionarily-divergent eukaryotes including the parasitic trypanosomatids, which are responsible for sleeping sickness (*Trypanosoma brucei*), Chagas disease (*Trypanosoma cruzi*), and leishmaniasis (*Leishmania* spp.) (Berriman et al., 2005; El-Sayed et al., 2005; Ivens et al., 2005). Using a yellow fluorescent protein (YFP)-tagging screen and mass spectrometry of co-purifying proteins, we previously identified 20 kinetochore proteins (KKT1–20) that localize to kinetochores in *T. brucei* (Akiyoshi and Gull, 2014; Nerusheva and Akiyoshi, 2016). These proteins have no obvious orthologs outside of kinetoplastids. More recently, KKT-interacting protein 1 (KKIP1) was identified as a kinetochore protein distantly related to Ndc80/Nuf2 based on similarity in the coiled-coil regions (D’Archivio and Wickstead, 2017). However, KKIP1 does not appear to have a CH domain, which is vital for the microtubule-binding activity of the Ndc80 complex. It is therefore unclear whether KKIP1 is a *bona fide* Ndc80/Nuf2-like protein. Affinity-purification of KKIP1 from cross-linked cells led to the identification of six additional proteins KKIP2–7 that localize to the kinetochore area during mitosis (D’Archivio and Wickstead, 2017). So far, very little is known about the function of these KKT and KKIP proteins.

The presence of putative DNA-binding motifs in KKT2 and KKT3 implies that these two proteins likely bind DNA (Akiyoshi and Gull, 2014). In contrast, none of the KKT or KKIP proteins has significant sequence similarity to the microtubule-binding domains present in canonical kinetochore proteins or microtubule-associated proteins, such as the CH domain (Ndc80/Nuf2, EB1) (Hayashi and Ikura, 2003; Wei et al., 2007; Ciferri et al., 2008), winged-helix domain (Ska1) (Jeyaprakash et al., 2012; Schmidt et al., 2012), TOG domain (XMAP215) (Slep, 2010; Al-Bassam and Chang, 2011), or spectrin fold (PRC1) (Subramanian et al., 2010). It therefore remains unknown which of the kinetochore proteins in kinetoplastids may bind microtubules.

It has been proposed that kinetoplastids represent one of the earliest-branching eukaryotes based on a number of unique ultrastructure and molecular features (Cavalier-Smith, 2010, 2013; Akiyoshi, 2016). Understanding how the kinetoplastid kinetochore mediates interactions with dynamic microtubules could therefore provide important insights into the evolutionary origins of kinetochore proteins as well as fundamental principles of kinetochore-microtubule coupling in eukaryotes.

Furthermore, because the molecular basis of microtubule interaction is likely distinct from other species, the kinetochores of kinetoplastids could be targeted to specifically kill those parasites that cause devastating diseases. In this study, we report the identification of the first microtubule-binding kinetochore component in *T. brucei*.

### KKT4 localizes at spindle microtubules in addition to kinetochores

In our previous analysis of endogenously YFP-tagged KKT proteins, we noticed that KKT4 localizes not only to kinetochores but also near the poles of the metaphase spindle (Akiyoshi and Gull, 2014). To further examine its cellular localization pattern, we used additional markers on kinetochores (KKT2) and on spindle microtubules (MAP103) (Hayashi and Akiyoshi, 2017) (Figure 1). These images confirmed that KKT4 has non-kinetochore signal that co-localizes with a spindle marker, especially in metaphase cells. Interestingly, some microtubule-binding kinetochore proteins (such as the Dam1 and Ska1 complexes) also decorate spindle microtubules (Cheeseman et al., 2001; Hanisch et al., 2006). It is therefore conceivable that KKT4 is located closely to microtubules, possibly making direct contacts.

**Figure 1.**
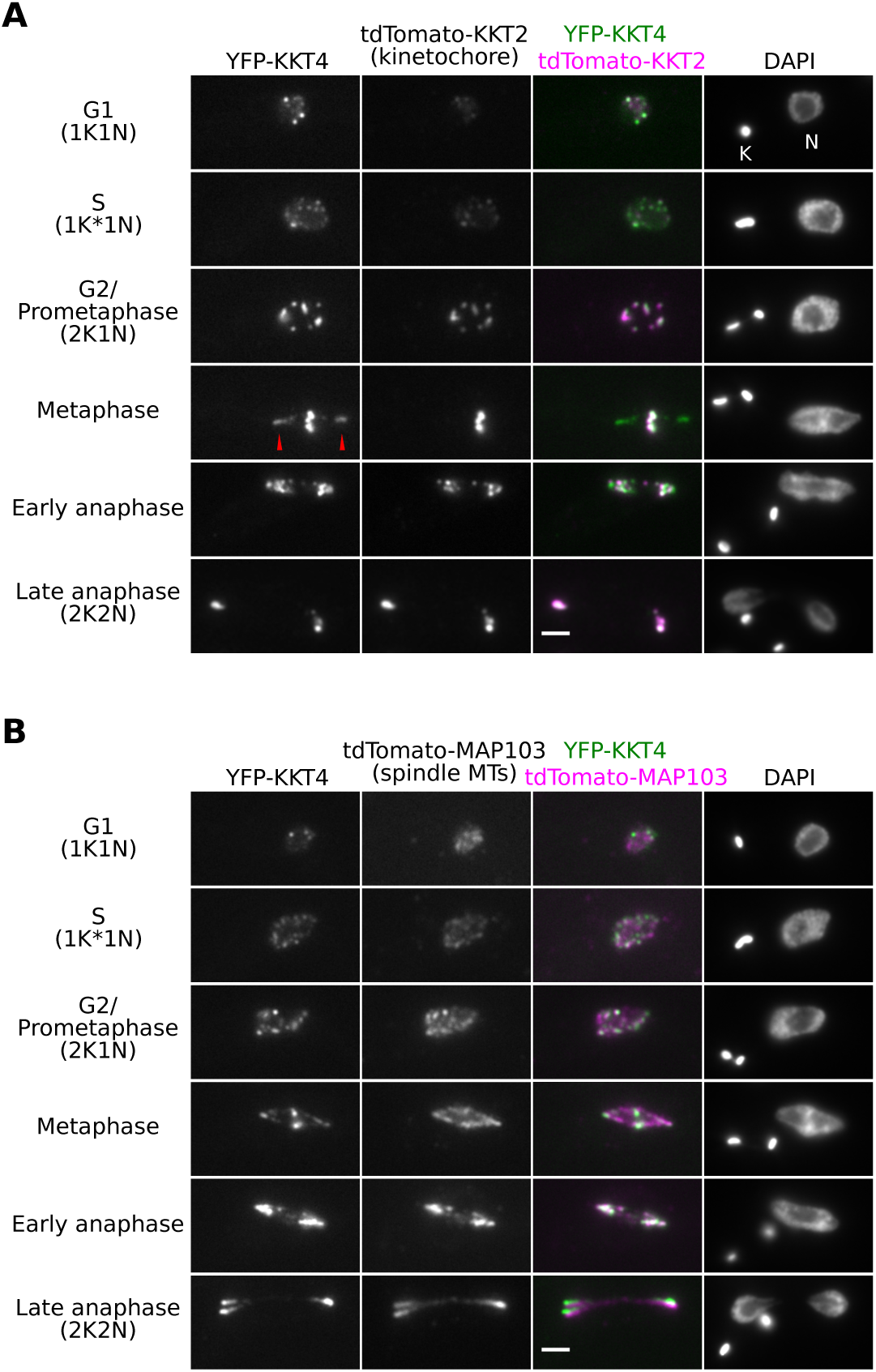
KKT4 localizes at spindle microtubules in addition to kinetochores. (A) Examples of cells expressing YFP-KKT4 and the kinetochore marker, tdTomato-KKT2, at indicated cell-cycle stages. Note that KKT4 has additional signals that do not co-localize with kinetochores. These additional, non-kinetochore KKT4 signals are especially prominent in metaphase (red arrowheads). K and N indicate the kinetoplast and nucleus, respectively. These organelles have distinct replication and segregation timings and serve as good cell-cycle markers (Woodward and Gull, 1990; Siegel et al., 2008). K* denotes an elongated kinetoplast and indicates that the nucleus is in S phase. (B) Examples of cells expressing YFP-KKT4 and the spindle microtubule marker, tdTomato-MAP103, showing that the non-kinetochore KKT4 signals partially co-localize with spindle microtubules. Bars, 2 µm.

### Depletion of KKT4 leads to a chromosome segregation defect

To examine the importance of KKT4 for chromosome segregation in vivo, we performed RNAi-mediated knockdown analysis. Upon induction of KKT4 RNAi, YFP-KKT4 signal was reduced at day 2 and growth retardation was observed by day 3 (Figure 2A, 2B). Chromosome segregation fidelity was then examined by monitoring the position of kinetochores in anaphase cells. We found that KKT4-depleted cells often had multiple lagging kinetochores (Figure 2B, 2C), showing that KKT4 is essential for proper chromosome segregation in vivo.

**Figure 2.**
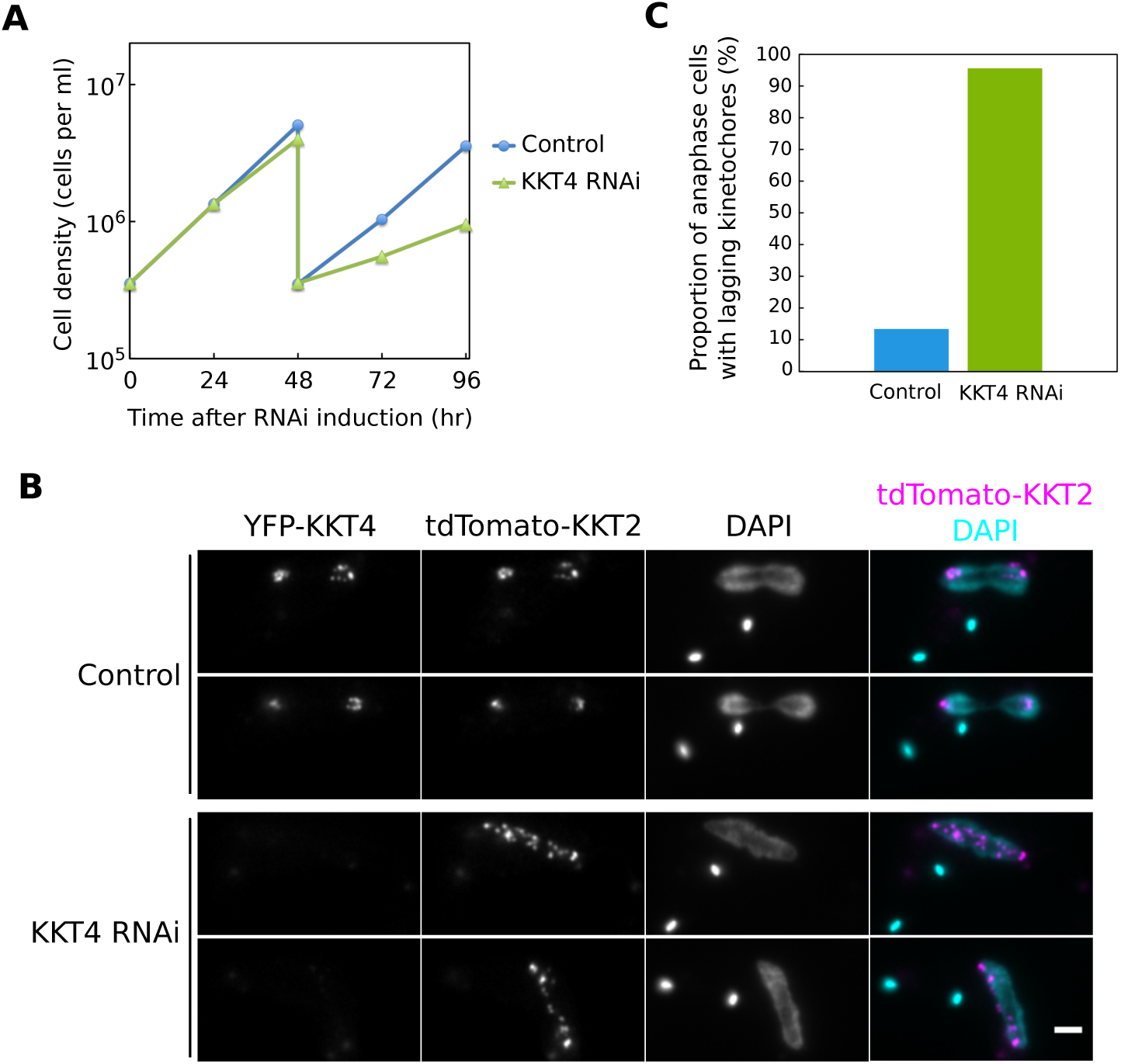
KKT4 is essential for accurate chromosome segregation. (A) RNAi-mediated knockdown of KKT4 leads to cell growth defect. Control is un-induced cell culture. Similar results were obtained using two independent RNAi constructs targeting different regions of the KKT4 transcript. (B) Cells expressing YFP-KKT4 and tdTomato-KKT2 were fixed at 48 hr after induction, showing a number of lagging kinetochores in anaphase. Note that YFP-KKT4 signal was reduced by RNAi. Bar, 2 µm. (C) Quantification of anaphase cells with lagging kinetochores at 48 hr post-induction (*n* > 100 each).

### KKT4 binds and diffuses along the microtubule lattice

To test whether KKT4 has direct microtubule-binding activity, we purified the full-length KKT4 protein fused with an N-terminal fluorescence and affinity tag, SNAP-6HIS-3FLAG. The protein was labeled during purification with a ^549^SNAP dye, and eluted from beads using FLAG peptides (Figure S1). We first tested whether the KKT4 protein binds taxol-stabilized microtubules using <u>t</u>otal <u>i</u>nternal <u>r</u>eflection <u>f</u>luorescence (TIRF) microscopy. Indeed, individual KKT4 particles bound transiently to coverslip-tethered microtubules, with an average residence time of 5.4 ± 0.5 s, and diffused along the filaments with a diffusion coefficient of 0.0071 ± 0.0003 µm^2^ s^−1^ (Figure 3). This lattice diffusion is highly reminiscent of the behavior of major microtubule-binding kinetochore proteins from other organisms, such as the human and yeast Ndc80 complexes (Powers et al., 2009), the Dam1 complex (Westermann et al., 2006), and the Ska complex (Schmidt et al., 2012).

**Figure 3.**
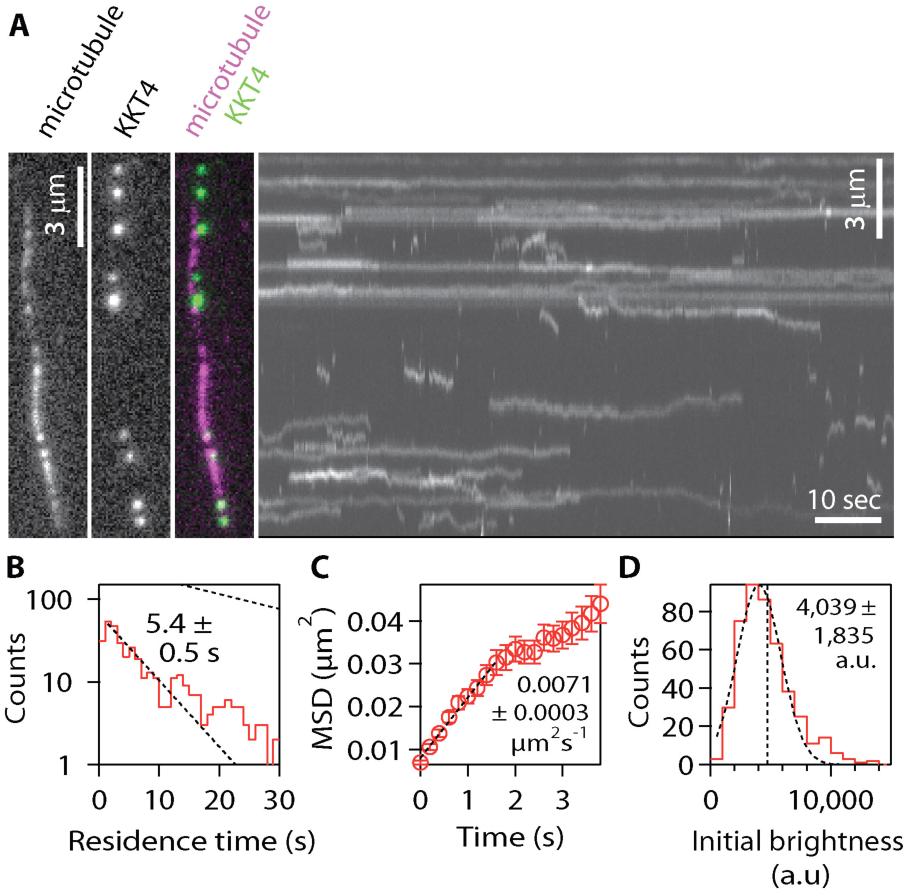
KKT4 binds and diffuses on microtubules. (A) Wild-type, fluorescent-tagged KKT4 particles (green) decorating a taxol-stabilized microtubule (magenta). (Left) Two-color fluorescence image. (Right) Corresponding kymograph showing diffusion of the KKT4 particles on the microtubule lattice. (B) Distribution of residence times on microtubules for wild-type KKT4 particles. Lower dotted line shows exponential fit used to determine average residence time (*n* = 452 binding events on 48 microtubules). Upper dotted line shows exponential distribution of bleach times for single fluorescent-tagged KKT4 particles, corresponding to an average of *τ*_*bleach*_ = 25 ± 1 s (*n* = 732 bleach events). (C) Mean-squared displacement (MSD) of wild-type KKT4 particles plotted against time. Dotted line shows linear fit used to determine diffusion coefficient (*n* = 452 events). (D) Distribution of initial brightness values for wild-type KKT4 particles diffusing on taxol-stabilized microtubules. Data are fitted by a Gaussian (dashed black curve) corresponding to a population with a unitary brightness of 4,039 ± 1,835 a.u. The vertical dashed line corresponds to the average brightness, 4,710 ± 2,323 a.u. (mean ± SEM; *n* = 452 particles).

Kinetochores of more commonly studied organisms contain arrays of microtubule-binding elements. The avidity of such arrays is believed to be crucial for maintaining persistent attachments to microtubule tips that are continuously assembling and disassembling (Hill, 1985; Joglekar et al., 2006; Powers et al., 2009; Alushin et al., 2010). To estimate the number of KKT4 molecules per microtubule-bound particle, we measured their brightness relative to that of an individual ^549^SNAP fluorophore (which was quantified from photobleaching experiments). These data suggest that, under the conditions of our assay, KKT4 binds the microtubule lattice in monomeric form (Figure 3D). Together, our data show that individual KKT4 molecules can bind microtubules directly and exhibit lattice diffusion, both of which are properties shared by core microtubule-binding kinetochore elements found in other organisms.

### Identification of basic residues important for KKT4 microtubule-binding activity

We next aimed to define the microtubule-binding domain within KKT4. The protein has several features conserved among kinetoplastids, including N-terminal predicted alpha helices (residues 2–30), predicted coiled-coils (121–225), a block of basic residues (326–340, predicted isoelectric point 11.0), and a C-terminal BRCT (BRCA1 C terminus (BRCT)) domain (462–645), which has been found in many DNA damage-response proteins and typically functions as a phosphorylation-dependent protein-protein interaction domain (Reinhardt and Yaffe, 2013) (Figure 4A and Figure S2). We initially purified four truncated forms of KKT4 and tested them in microtubule co-sedimentation assays. Only the central fragment with the predicted coiled-coils and basic block, KKT4^115–343^, co-sedimented robustly with taxol-stabilized microtubules, while the other three constructs did not (Figure 4B). To further dissect the protein, we created two shorter constructs, KKT4^115–174^ and KKT4^168–343^. Of these, only KKT4^115–174^ co-sedimented with microtubules (Figure 4C). These observations suggest that KKT4^115–174^ represents a minimal microtubule-binding domain. However, we note that KKT4^115–174^ co-sedimented less efficiently than KKT4^115–343^, implying that a feature within residues 175–343 enhances the activity of the minimal microtubule-binding domain.

**Figure 4.**
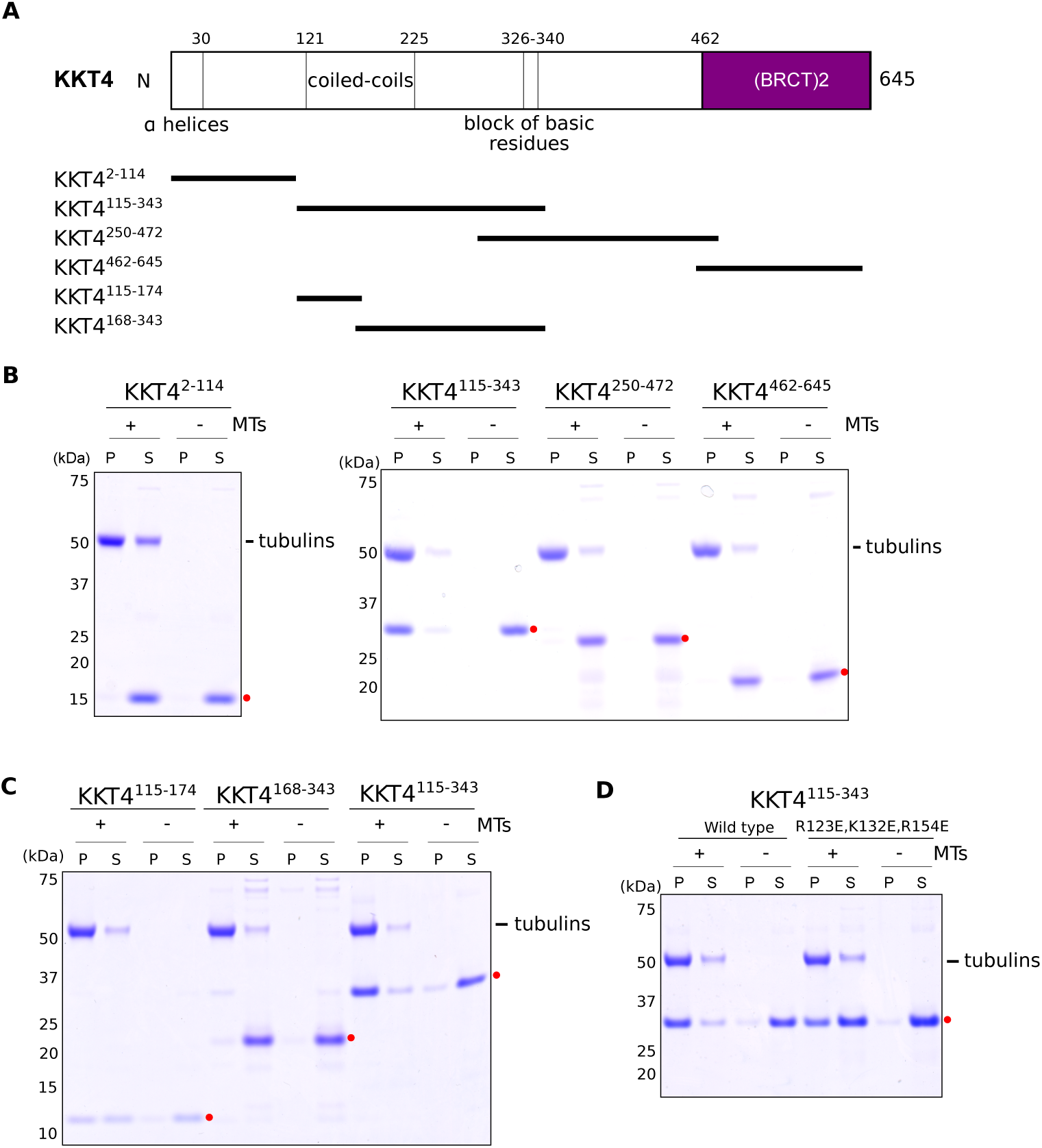
KKT4^115–174^ is sufficient for microtubule-binding. (A) Schematic of the KKT4 protein sequence. Several features conserved among kinetoplastids are shown, with the truncated proteins used in Figure 4B and 4C indicated below. (B) Microtubule sedimentation assays of KKT4 fragments showing that KKT4^115–343^ can bind microtubules, whereas KKT4^2-114^, KKT4^250-472^, and KKT4^462-645^ do not. P and S stand for pellet and supernatant, respectively. Red dots indicate KKT4 fragments tested in the assay. (C) Microtubule sedimentation assays of KKT4 fragments showing that KKT4^115–174^ can bind microtubules, albeit to a lesser extent than KKT4^115–343^. (D) The charge-reversal mutant of KKT4 has reduced microtubule-binding activity.

Due to its positively charged residues, the predicted isoelectric point of KKT4^115–174^ is 9.8 (while that of KKT4^115–343^ is 9.7). Because microtubule binding is often mediated by electrostatic charges (Ciferri et al., 2008; Schmidt et al., 2012), we replaced three basic residues within the KKT4^115–343^ construct with acidic residues (R123E, K132E, and R154E). This charge-reversal mutant sedimented with taxol-stabilized microtubules less efficiently than wild-type KKT4^115–343^ (Figure 4D), indicating that its microtubule-binding activity was compromised. Likewise, in TIRF experiments (Figure 5A), the average residence times on taxol-stabilized microtubules for individual mutant KKT4 particles, carrying the same set of charge-reversal mutations, were ~2-fold shorter compared to wild-type KKT4 (Figure 5B) and their lattice diffusion was ~2-fold faster (Figure 5C). Similar changes in single-particle residence time and lattice diffusion occur when mutations or secondary modifications (phosphorylations) are introduced into the microtubule-binding domains of kinetochore proteins from other organisms (Gestaut et al., 2008; Umbreit et al., 2012; Zaytsev et al., 2014). Thus, our observations suggest that the positively charged residues, R123, K132, and R154, make an important contribution to the microtubule-binding activity of KKT4.

**Figure 5.**
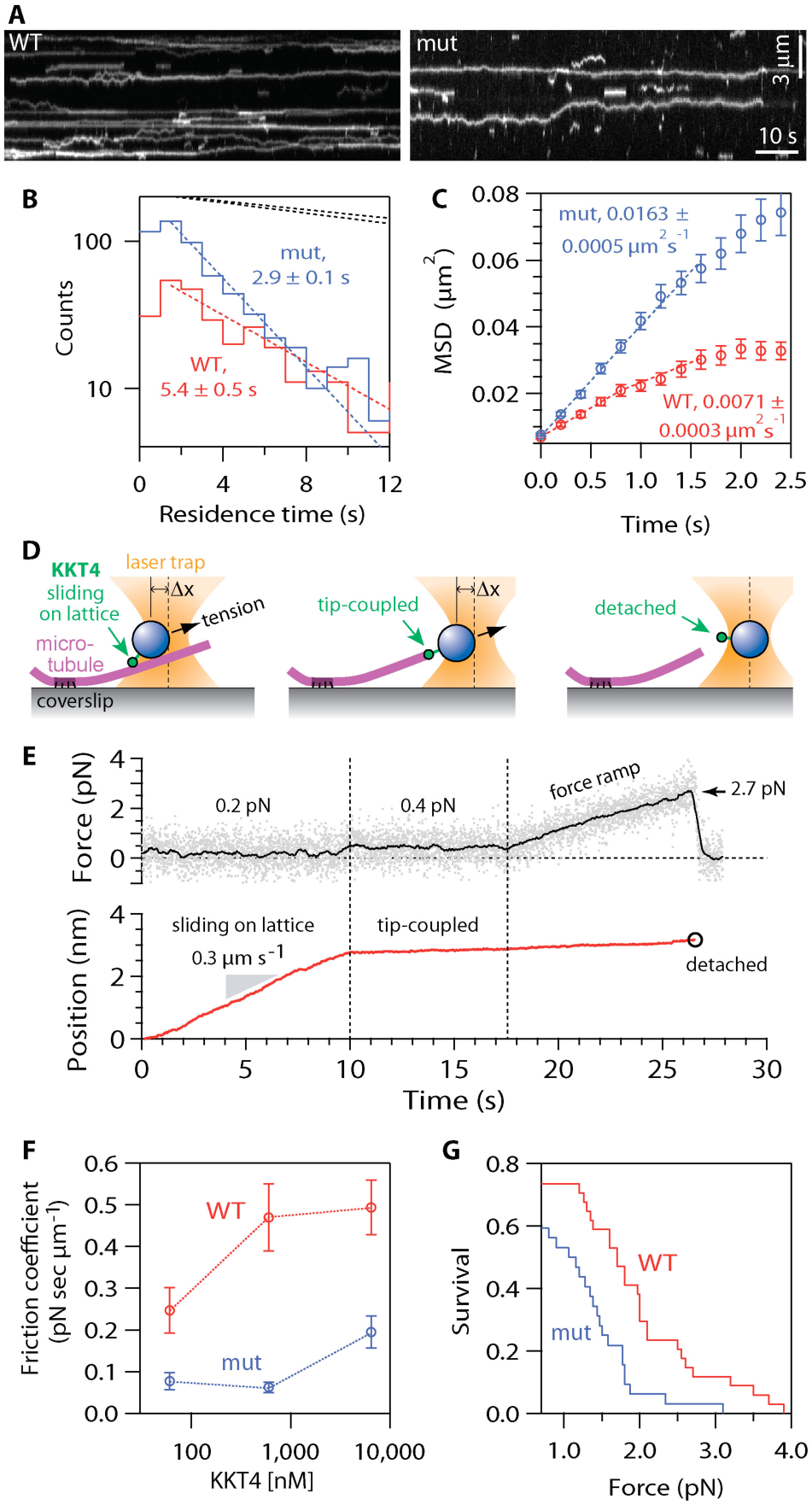
Charge-reversal mutant KKT4 binds more weakly. (A) Kymographs showing binding and diffusion of full-length, wild-type KKT4 (WT) and charge-reversal mutant KKT4 (mut) on taxol-stabilized microtubules. (B) Distributions of residence times on microtubules for wild-type KKT4 (red) and charge-reversal mutant KKT4 (blue). Corresponding dotted lines show exponential fits used to determine average residence times (*n* > 452 binding events on > 48 microtubules). Upper dotted lines show exponential bleach-time distributions for single wild-type and mutant KKT4 particles, corresponding to average bleach times of *τ*_*bleach*_ = 25 ± 1 s and *τ*_*bleach*_ = 30 ± 1 s, respectively. Wild-type data are recopied from Figure 3 for comparison. (C) Mean-squared displacement (MSD) of wild-type KKT4 (red) and mutant KKT4 (blue) particles plotted against time. Dotted lines show linear fits used to determine diffusion coefficients (*n* > 452 particles). Wild-type data are recopied from Figure 3 for comparison. (D) Schematic of laser trap assay used to measure friction coefficients and rupture strengths for KKT4^115-343^-decorated beads. (E) Example record showing trap force and bead displacement versus time. (F) Friction coefficients for wild-type KKT4^115-343^ (red) and charge-reversal mutant KKT4^115-343^ (blue) at indicated concentrations (mean ± SEM; *n* = 13 to 47 events). All individual friction coefficient values are given in Table S1. (G) Attachment survival probability versus force for wild-type KKT4^115-343^ (red) and charge-reversal mutant KKT4^115-343^ (blue; *n* = 33 and 35 events, respectively). All individual rupture force values are given in Table S1.

### KKT4 forms load-bearing attachments to dynamic microtubule tips

Kinetochore-microtubule attachments in vivo must bear piconewton-scale loads, especially during prometaphase, when bioriented chromosomes are subjected to tensile forces from opposing microtubules (Nicklas, 1988; Chacón et al., 2014; Ye et al., 2016). To measure the load-bearing capacity of KKT4-based attachments, we applied a computer-controlled laser trap, adapting assays developed previously for the study of kinetochore components from other organisms (Asbury et al., 2006; Powers et al., 2009; Franck et al., 2010). Polystyrene microbeads were decorated with purified KKT4^115–343^ protein and then attached to individual dynamic microtubules growing from coverslip-anchored seeds. Initially, the beads were attached laterally to the sides of the microtubules, in-between the coverslip anchor and the growing filament tip (Figure 5D, left). This arrangement mimics the in vivo situation where kinetochores initially attach laterally to spindle microtubules (Hayden et al., 1990; Tanaka et al., 2005). Constant tensile forces of ~1 pN, applied using feedback-control toward the plus end, caused the laterally attached beads to slide along the filament lattice, usually without detachment (Figure 5E), similar to the lateral sliding behavior seen previously in vitro with other kinetochore components (Asbury et al., 2006; Powers et al., 2009). Beads decorated with the charge-reversal mutant KKT4^115-343^ slid more easily along the microtubule than those decorated with the same concentrations of wild-type KKT4^115-343^. To quantify this difference, we calculated friction coefficients (Bormuth et al., 2009), dividing the applied force by the sliding speeds (Figure 5F and Table S1, see materials and methods). The average friction coefficient for charge-reversal mutant-coated beads was substantially lower than that for wild-type across a range of concentrations, indicating a weaker interaction of the mutant with the lattice. Once a sliding bead reached the plus end of the microtubule, its movement abruptly slowed and was thereafter governed by the speed of microtubule assembly (Figure 5D, middle). This ‘tip-coupled’ arrangement mimics the situation in vivo when kinetochores maintain load-bearing attachments to growing plus ends (Inoué and Salmon, 1995). To measure the strength of tip-coupling, we increased the tension gradually (at 0.25 pN s^−1^, under feedback-control) until the attachment ruptured (Figure 5D, right). Tip attachments formed by wild-type KKT4^115–343^ ruptured at significantly higher forces compared to the charge-reversal mutant-based attachments (Figure 5G and Table S1). These observations establish that KKT4-based couplers can support significant loads and they confirm that the strengths of both lateral and end-on attachments to dynamic microtubules depends on the three basic residues (R123, K132, R154).

### KKT4 tracks with dynamic microtubule tips and harnesses tip disassembly to produce force

The forces that move chromosomes during prometaphase, metaphase, and anaphase in vivo are generated in part via the tracking of kinetochores with dynamic microtubule tips (Inoué and Salmon, 1995). By maintaining a persistent, load-bearing attachment to a disassembling microtubule tip, a kinetochore harnesses energy released from the tip to produce mechanical work (i.e., force acting through distance) (Hill, 1985; Koshland et al., 1988; Coue et al., 1991). We found that purified KKT4 alone can recapitulate this tip-coupling activity. Using TIRF microscopy, we observed individual particles of fluorescently-labelled full-length KKT4 bound along dynamic microtubules and then induced disassembly of the filaments by washing out free tubulin. When disassembling tips encountered KKT4 particles, the particles began tracking with the tips and were usually carried all the way to the coverslip-anchored seed (Figure 6A and Movie S1). Likewise, microbeads decorated with KKT4^115–343^ tracked persistently with disassembling microtubule tips in the absence of externally applied force (Figure 6B and Movie S2). To test the load-bearing capacity, we applied constant forces to tip-tracking beads using feedback-control. We found that KKT4^115–343^ reliably tracked with both assembling and disassembling microtubule tips under constant forces of ~1 pN (Figure 6C). This load-bearing capacity is comparable to other microtubule-binding kinetochore components such as the Dam1 and Ndc80 complexes (Asbury et al., 2006; Powers et al., 2009), suggesting that KKT4 is directly involved in chromosomes segregation.

**Figure 6.**
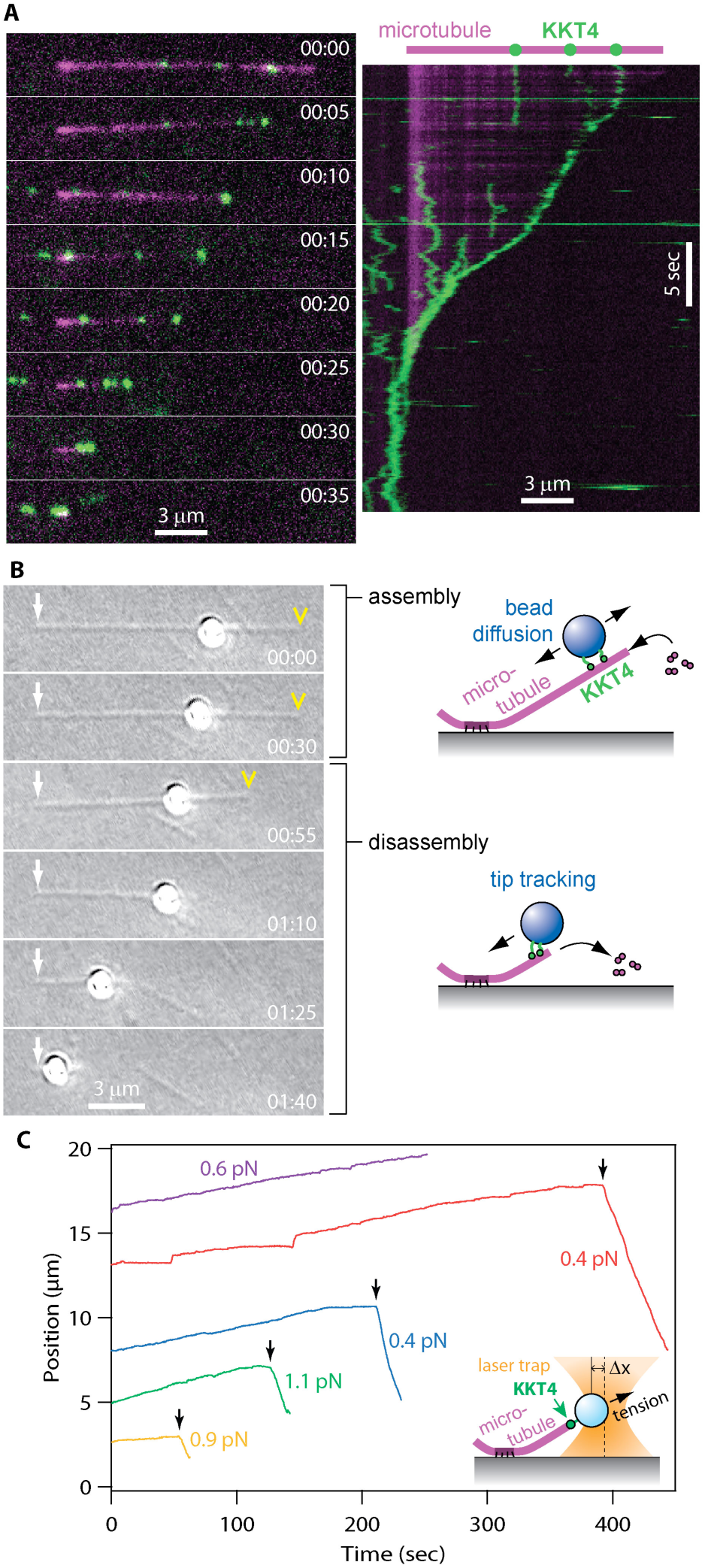
KKT4 tracks with dynamic microtubule tips. (A) Selected frames (left) and kymograph (right) from Movie S1 showing wild-type KKT4 (green) tracking with the disassembling tip of a microtubule (magenta). Elapsed times are in min:sec. Individual KKT4 particles can also be seen diffusing on the microtubule lattice. (B) Selected frames from Movie S2 showing a wild-type KKT4^115–343^-coated bead diffusing on the microtubule lattice and then tracking with a disassembling tip. White arrows indicate the coverslip-anchored portion of the microtubule seed. Yellow arrowheads indicate the microtubule tip. Elapsed times are in min:sec. (C) Records of bead position versus time during continuous application of tensile force. Increasing position represents assembly-coupled movement in the direction of applied force, away from the coverslip-anchored seed (e.g., red trace, < 400 s). Decreasing position represents disassembly-driven motion against the applied force (e.g., red trace, > 400 s). Arrows (↓) indicate ‘catastrophe’ events, when the microtubule tip switched spontaneously from assembly into disassembly. For clarity, the records are offset vertically by an arbitrary amount. (Inset) Schematic of laser trap assay.

## Conclusions

In this study, we report the identification of KKT4 as the first microtubule-binding kinetochore protein in *Trypanosoma brucei*. Its microtubule-binding domain lacks significant similarity to any other microtubule-binding protein (Akiyoshi and Gull, 2014). Furthermore, there is no kinetochore protein known in other eukaryotes that has a BRCT domain (Musacchio and Desai, 2017). These results suggest that KKT4 is a unique kinetochore protein that evolved from an ancestral BRCT-domain containing protein to interact with microtubules in kinetoplastids. Lack of similarity to other microtubule-binding proteins raises a possibility that KKT4 may interact with microtubules in a distinct manner. Structural analysis of KKT4 will be important to elucidate the mode of microtubule-binding by this protein.

During the purification of full length KKT4 protein from insect cells, we noticed co-purification of insect DNA, which could be removed by a heparin affinity chromatography column. This observation suggests that KKT4 has DNA-binding activities at least in vitro. Although we do not yet know its physiological significance, the fact that KKT4 is constitutively localized at the kinetochore supports the possibility that KKT4 may also interact with DNA in vivo, which warrants further studies in the future. In prokaryotes, plasmid DNA segregation is mediated by a DNA-binding protein that also interacts with polymers (e.g. ParR protein associates with *par*C DNA and filament forming ParM protein) (Garner et al., 2007; Gerdes et al., 2010). We speculate that KKT4 could in principle bridge both DNA and microtubules like prokaryotic segregation proteins. Furthermore, our previous finding that KKT4 co-purified with APC/C subunits (Akiyoshi and Gull, 2014) also raises a possibility that KKT4 may directly regulate the timing of anaphase onset upon establishment of proper microtubule attachments. Further characterization of KKT4 is key to better understanding how the unconventional kinetoplastid kinetochore performs conserved kinetochore functions.

## Materials and Methods

### Trypanosome cells

All trypanosome cell lines used in this study were derived from *T. brucei* SmOxP927 procyclic form cells (TREU 927/4 expressing T7 RNA polymerase and the tetracycline repressor to allow inducible expression) (Poon et al., 2012) and are listed in Table S2. Plasmids and primers used in this study are listed in Table S3 and S4, respectively. Cells were grown at 28ºC in SDM-79 medium supplemented with 10% (v/v) heat-inactivated fetal calf serum (Brun and Schönenberger, 1979). Cell growth was monitored using a CASY cell counter and analyzer system (Roche). RNAi was induced with doxycycline at a final concentration of 0.1 µg mL^−1^. Endogenous YFP and tdTomato tagging was performed using the pEnT5-Y vector (Kelly et al., 2007) and pBA148 (Akiyoshi and Gull, 2014), respectively. pBA215 (TY-tdTomato-MAP103 tagging vector) was made by sub-cloning a *Xba*I/*Bam*HI fragment of pBA31 (Hayashi and Akiyoshi, 2017) into pBA148. For generation of an inducible KKT4 stem-loop RNAi construct, the following DNA fragment was synthesized by GeneArt and cloned into the pBA310 inducible expression vector (Akiyoshi and Gull, 2014) using *Hin*dIII and *Bam*HI sites: a *Hin*dIII site, 428 bp fragment targeting KKT4 3’UTR (starting at 63 bp downstream of the KKT4 stop codon), stuffer sequence (AAAGGCGGACCCTCATTTCTAAGTACGGTCAGGTGTCGTAGCACTGCATT GAATTCGATTGCCATTCTCCGAGTGTTTTAGCGTGACGGCCGCAGGGGTCCCATAA), reversed orientation of the 428 bp fragment, and *Bam*HI site. Similar results were obtained using two independent RNAi constructs targeting the coding sequence of KKT4 (23–455 and 946–1348) (data not shown). Plasmids linearized by *Not*I were transfected to trypanosomes by electroporation into an endogenous locus (pEnT5-Y and pBA148 derivatives) or 177 bp repeats on minichromosomes (pBA310 derivatives). Transfected cells were selected by the addition of 25 µg mL^−1^ hygromycin (pEnT5-Y derivatives), 10 µg mL^−1^ blasticidin (pBA148 derivatives), or 5 µg mL^−1^ phleomycin (pBA310 derivatives). Microscopy was performed as previously described (Nerusheva and Akiyoshi, 2016).

### Protein expression and purification from *E. coli* and insect cells

DNA fragments encoding truncated KKT4 were amplified from genomic DNA and cloned into the pNIC28-BsaI expression vector (gift of SGC) using a ligation-independent cloning method (Gileadi et al., 2008). Truncated KKT4 proteins fused with N-terminal 6HIS tag and a TEV protease cleavage site were expressed in *Escherichia coli* BL21(DE3) cells. Cells were grown in 2xTY media at 37ºC to an OD600 of ~0.6. Protein expression was induced by 0.1 mM IPTG and incubated overnight at 16ºC. Cells were pelleted at 3400 g at room temperature, and cell pellet was frozen in liquid nitrogen and stored in −80ºC. We resuspended cells in lysis buffer (50 mM sodium phosphate pH 7.5, 500 mM NaCl, 10% glycerol) supplemented with protease inhibitors (Leupeptin, Pepstatin, E-64, 20 µg mL^−1^ each, and 2 mM benzamidine) and 1 mM TCEP, and sonicated cells on ice. Lysed cells were spun at 48,000 g at 4ºC for 25 min. Supernatant was incubated with Talon beads (Takara Clontech) for 1 hr at 4ºC. We extensively washed the beads with lysis buffer, and eluted proteins with elution buffer (50 mM sodium phosphate pH 7.5, 500 mM NaCl, 10% glycerol, 250 mM imidazole). Buffer was exchanged into BRB80 (80 mM PIPES-KOH pH 6.9, 1 mM EGTA, 1 mM MgCl_2_) with 100 mM KCl using a PD MiniTrap G-25 column (GE). We then concentrated the sample by 3 kDa or 10 kDa MW Amicon concentrator (Millipore), flash froze aliquots in liquid nitrogen, and stored in −80ºC.

Synthetic DNA (GeneArt) encoding full-length KKT4 (codon-optimized for expression in insect cells) fused with an N-terminal SNAP-6HIS-3FLAG tag was cloned into the pACEBac2 vector (Geneva Biotech) (Bieniossek et al., 2012). Bacmid was purified from DH10EmBacY *E. coli* cells using PureLink HiPure Plasmid Miniprep Kit (Thermo Fisher), and used to transfect Sf9 cells using Cellfectin II transfection reagent (Thermo Fisher). Sf9 cells were grown in Sf-900 II SFM media (Thermo Fisher). Baculovirus was amplified through three rounds of amplification. Typically, 500 ml culture of Sf9 cells at 1–1.2 million cells mL^−1^ was infected with P3 baculovirus for ~64 hours before harvesting. Subsequent steps were carried out at 4ºC. Cells were pelleted at 700 g for 10 min, washed once with PBS, and resuspended in 10 ml of BH0.25 (25 mM HEPES pH 7.5, 2 mM MgCl_2_, 0.1 mM EDTA, 0.5 mM

EGTA, 10% glycerol and 250 mM NaCl) supplemented with 2x protease inhibitors (Leupeptin, Pepstatin, E-64, 20 µg mL^−1^ each, and 0.4 mM PMSF) and 0.01 µg mL^−1^ DNase I. We added 0.2% NP-40 (Igepal CA-630) to the sample and lysed cells by a dounce homogenizer (3 rounds of 10 strokes with 5 min break in between). The sample was diluted by adding 12.5 ml of BH0.25 with protease inhibitors, and centrifuged for 30 min at 45,000 g. The supernatant was incubated with 2 ml Anti-FLAG M2 affinity gel (Sigma) for 3 hours with constant rotation, followed by five washes with BH0.25 with 1x protease inhibitors and 2 mM DTT (20 ml each). Beads were incubated with 2.5 ml of 20 µM ^549^SNAP dye (New England Biolabs) for 30 min at room temperature, and subsequently washed twice with BH0.25 with 1x protease inhibitors and DTT, and twice with BH0.25 with 1x protease inhibitors. Proteins were eluted from the beads with gentle agitation of beads in 2 ml of BH0.25 containing 0.5 mg mL^−1^ 3FLAG peptide (Sigma) and 1x protease inhibitors. The sample was further purified using 1 ml HiTrap Heparin HP column pre-equilibrated with 5% of buffer B (buffer A: 20 mM HEPES pH 7.5 with 1 mM TCEP, buffer B: 20 mM HEPES pH 7.5, 1 M NaCl with 1 mM TCEP), and eluted with a linear gradient from 5% to 100% of buffer B. Fractions containing ^549^SNAP-tagged KKT4 were pooled (eluted at ~340 mM NaCl) and flash-frozen in liquid nitrogen and stored at - 80ºC before being used for TIRF or laser trap assays. The labeling was confirmed by an FLA 7000 scanner using the SHG532 laser and 0580 filter (GE). For analytical size exclusion chromatography, the sample was concentrated to ~24 µM by 10 kDa MW Amicon concentrator (Millipore), and loaded onto a Superose 6 increase 5/150 GL column (GE) equilibrated in gel filtration buffer (25 mM HEPES pH 7.5, 150 mM NaCl with 1 mM TCEP) on an ÄKTA pure 25 system. Protein concentration was determined by comparing the purified samples with BSA standards on SDS-PAGE gels as well as Protein Assay (Bio-rad).

### Co-sedimentation assay

Taxol-stabilized microtubules were prepared as follows. We mixed 2.5 µL of 100 µM porcine tubulin resuspended in BRB80 with 1 mM GTP (Cytoskeleton Inc), 1.25 µL of BRB80, 0.5 µL of 40 mM MgCl_2_, 0.5 µL of 10 mM GTP, and 0.25 µL of DMSO, and incubated for 20 min at 37ºC. Then we added 120 µL of BRB80 containing 12.5 µM taxol (Paclitaxel, Sigma) to the sample, and passed through a 27G1/2 needle once to prepare sheared microtubules (2 µM). For microtubule co-sedimentation assays, 20 µL of KKT4 fragments (at final concentration of 4 µM) and 20 µL of microtubules (final 1 µM) were mixed and incubated for 45 min at room temperature. For a no-microtubule control, we incubated KKT4 fragments with BRB80 with 12.5 µM taxol. The samples were spun at 20,000 g at room temperature for 10 min, and supernatant was taken. We added 40 µL of chilled BRB80 with 5 mM CaCl_2_ and put the samples on ice for 5 min to depolymerize microtubules. All samples were boiled for 3 min before analysis by SDS-PAGE gels stained with Coomassie Brilliant Blue R-250 (Bio-rad).

### TIRF binding assays

Recombinant, ^549^SNAP-tagged, full-length, wild-type KKT4 or the charge-reversal mutant KKT4 were used for all of the TIRF experiments. Flow channels were prepared using silanized coverslips as previously described (Gestaut et al., 2010; Kudalkar et al., 2016), coated with 1 mg mL^−1^ biotinylated BSA (Vector laboratories), and thoroughly washed with BRB80. Then 1 mg mL^−1^ avidin DN (Vector laboratories) was added and the chamber was washed with BRB80. Then biotinylated, taxol-stabilized microtubules were introduced, allowed to bind the coverslip surface, and then washed with a wash buffer of BRB80, 1 mM DTT, 10 µM Taxol, and 1 mg mL^−1^ k-casein. Finally, 1.5 nM ^549^SNAP-tagged KKT4 was added in a buffer of BRB80, 1 mg mL^−1^ k-casein, 1 mM DTT, 10 µM Taxol, 250 µg mL^−1^ glucose oxidase, 25 mM glucose, and 30 µg mL^−1^ catalase. The chamber was allowed to incubate for 5 min before imaging in a custom-built TIRF microscope (Deng and Asbury, 2017). Images were acquired at 5 frames sec^−1^ for 100 sec.

### TIRF depolymerization assays

Flow chambers were prepared and microtubules were attached using a similar method as described above for taxol-stabilized microtubule binding assays, but with minor modifications. First, 1 mg mL^−1^ biotinylated BSA (Vector Laboratories) was flowed in and incubated for 5 min. The chamber was then washed with BRB80 followed by an incubation with 1 mg mL^−1^ avidin DN (Vector Laboratories) for 5 min. The chamber was again washed, and GMPCPP-stabilized microtubule seeds (Hyman et al., 1991; Asbury et al., 2006) were flowed in and allowed to bind the surface for 5 min. The chamber was washed off excess seeds by introducing microtubule growth buffer, consisting of BRB80, 1 mM DTT, 1 mM GTP, and 1 mg mL^−1^ k-casein. Then, in the same growth buffer, 10 µM of free tubulin and 3.5 nM of wild-type KKT4 were introduced into the chamber. The chamber was incubated at 30°C for 10 min before imaging. Once it was clear that microtubule extensions of sufficient length had grown, their disassembly was promoted by exchanging buffer without tubulin, but with 3.5 nM KKT4.

### Brightness and Dwell Time Analysis

Custom TIRF analysis software was developed using Labview (National Instruments) as previously described (Gestaut et al., 2010; Asbury, 2016). This software generated kymographs of KKT4, which could be used to get the position and brightness over time. Histograms of KKT4 particle brightness and dwell time on microtubules, and plots of mean squared displacement versus time, were generated using Igor Pro (WaveMetrics).

### Laser trap assays

Recombinant 6HIS-KKT4^115–343^ was attached to 0.56 µm diameter streptavidin-coated polystyrene beads (Spherotech, Lake Forest, IL) using biotinylated His5 antibody (QIAGEN, Valencia, CA) as previously described (Franck et al., 2010). The amount of protein per bead was adjusted by incubating dilutions of 60, 600, and 6,500 nM, prepared in BRB80, 8 mg mL^−1^ BSA, and 1 mM DTT with a fixed concentration of beads (0.025% w/v) at 4ºC for 1 hour. Our laser trapping-based motility assay has been previously described (see, for example (Asbury et al., 2006; Powers et al., 2009; Franck et al., 2010)). Briefly, dynamic microtubule extensions were grown from coverslip-anchored GMPCPP-stabilized microtubule seeds in a buffer consisting of BRB80, 1 mg mL^−1^ κ-casein, 1 mM GTP, 250 µg mL^−1^ glucose oxidase, 25 mM glucose, 30 µg mL^−1^ catalase, 1 mM DTT and 6 µM (dimer) of purified bovine brain tubulin.

The laser trap has been previously described (Franck et al., 2010). Force feedback was implemented with custom LabView software. During clamping of the force, bead-trap separation was sampled at 40 kHz while stage position was updated at 50 Hz to maintain the desired load. Bead and stage position data were decimated to 200 Hz before storing to disk.

### Rupture force and friction coefficient measurements

Beads were prepared with either wild-type or charge-reversal mutant KKT4^115–343^ at dilutions of 60, 600 and 6,500 nM. Individual beads were attached to the microtubule lattice and preloaded with a constant tension of 0.8 ± 0.3 pN. To measure the lateral friction coefficient (γ) we measured the force at which the bead was dragged along the lattice (F) and divided by the velocity at which the piezo stage moved (v), γ = F v^−1^. Once the bead reached the microtubule tip, the laser trap was programmed to ramp the force at a defined rate (0.25 pN s^−1^) until the linkage ruptured. For the experiments in which tip-tracking beads were observed in the absence of laser trapping force (Figure 6B), the beads were coated with wild-type KKT4^115–343^ at 60 nM.

## Supplemental Information

Supplemental Information includes 2 movies, 3 figures, and 4 tables.

## Author contributions

A.L., H.H., M.E.B., C.L.A., and B.A. conceived the experiments. H.H. and B.A. generated and analyzed trypanosome cell lines. H.H., A.W., and P.L. purified KKT4 fragments and performed sedimentation assays. H.H. and B.A. purified and analyzed fluorescently labelled KKT4. A.L. and M.E.B. performed the fluorescence measurements. A.L. performed laser trap experiments. A.L., H.H., M.E.B., C.L.A., and B.A. prepared the manuscript.

## Acknowledgments

We thank Gabriele Marcianò, Midori Kanazawa, and Olga Nerusheva for comments on our manuscript. We also thank the Micron Oxford Advanced Bioimaging Unit. A.L. was supported by a Sackler Scholars Fellowship in Integrative Biophysics. H.H. was supported by a Uehara Memorial Foundation fellowship. M.E.B. was supported by an NIH Interdisciplinary Training Fellowship (T32CA080416). C.L.A. was supported by grants from the NIH (RO1GM079373, P01GM105537) and by a David and Lucile Packard Fellowship (2006-30521). B.A. was supported by a Sir Henry Dale Fellowship jointly funded by the Wellcome Trust and the Royal Society (grant number 098403/Z/12/Z), Wellcome-Beit Prize Fellowship (grant number 098403/Z/12/A), and the EMBO Young Investigator Program.

## Supplemental Materials for

This PDF file includes

- Caption for Movie S1, S2
- Supplemental Figure S1–3
- Caption for Supplemental Table S1
- Supplemental Tables S2–4

Other Supplemental Materials for this manuscript include the following

- Movie S1 and S2 (AVI)
- Table S1 (Excel)

### Supplemental Movies

**Movie S1**. **Wild-type KKT4 tracks with a disassembling microtubule tip.** TIRF movie of wild-type, ^549^SNAP-tagged KKT4 (green) tracking with a depolymerizing, Alexa^488^-labeled microtubule (magenta). KKT4 concentration was 3.5 nM. To induce microtubule depolymerization, free tubulin (10 µM) was washed out from the chamber. The movie corresponds to the event in Figure 6A. It was recorded at 5 frames s^−1^.

**Movie S2**. **A bead decorated with wild-type KKT4^115–343^ diffuses on the microtubule lattice and then undergoes disassembly-driven motion.** Microtubule depolymerization drives the movement of the bead towards the microtubule minus end. The contrast-enhanced movie corresponds to the event in Figure 6B, which was recorded at 30 frames s^−1^. The bead was decorated at 60 nM of KKT4.

**Figure S1.**
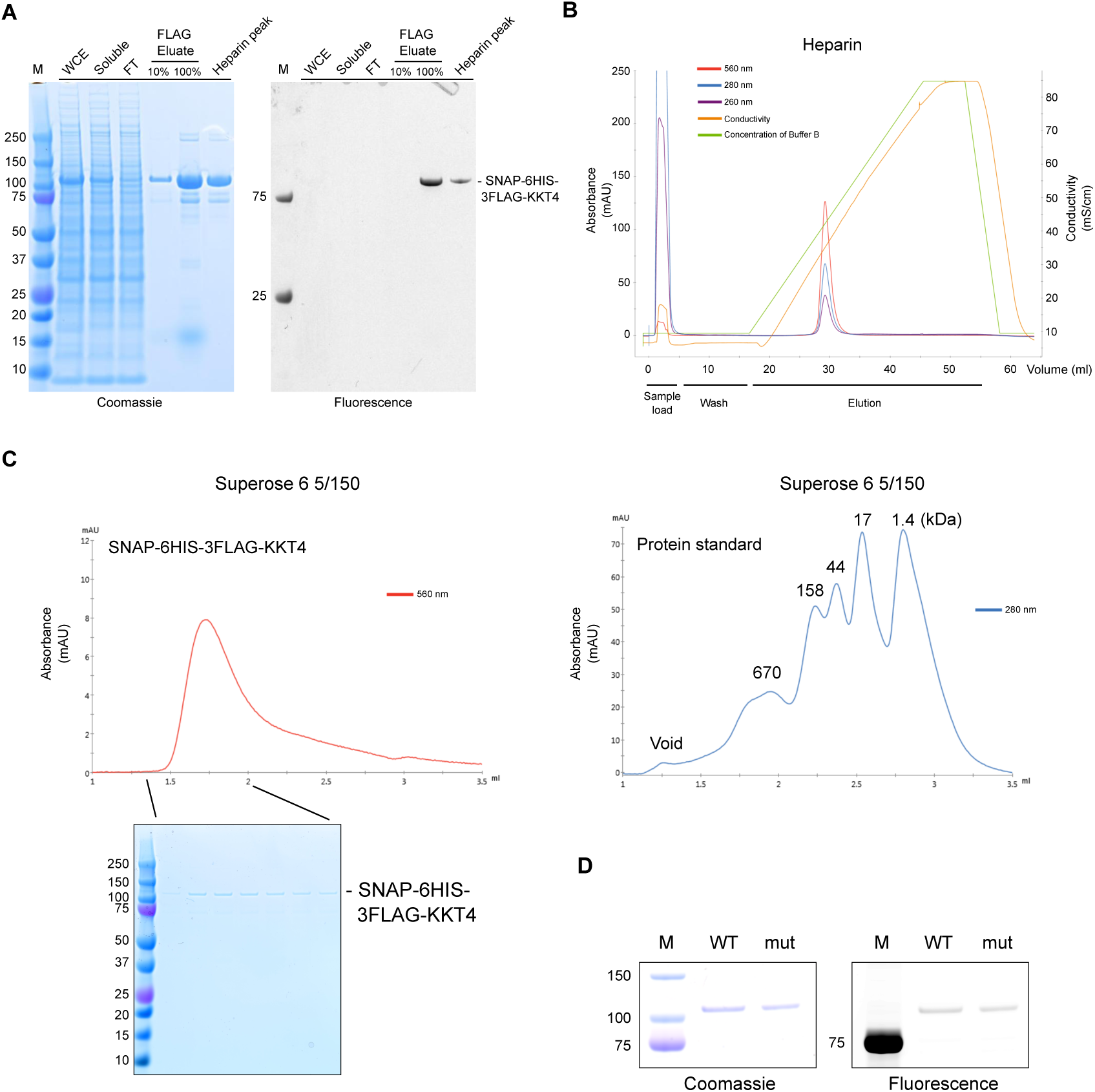
Purification and characterization of full length KKT4 used for TIRF assays. (A) SNAP-6HIS-3FLAG-KKT4 was expressed and purified from insect cells. The protein was labeled with ^549^SNAP during purification. Left: Coomassie-stained SDS-PAGE gel. Right: fluorescence scan of the same gel. WCE and FT stand for whole cell extract and flow through, respectively. (B) Heparin chromatography of ^549^SNAP-6HIS-3FLAG-KKT4 sample to get rid of 3FLAG peptides and DNA contamination. (C) Size-exclusion chromatography of ^549^SNAP-6HIS-3FLAG-KKT4, showing that it migrates as a single peak in the Superose 6 5/150 column (left). Protein standard is shown on the right. (D) Purification of WT and charge-reversal mutant (mut: R123E, K132E, R154E) of ^549^ SNAP-6HIS-3FLAG-KKT4.

**Figure S2.**
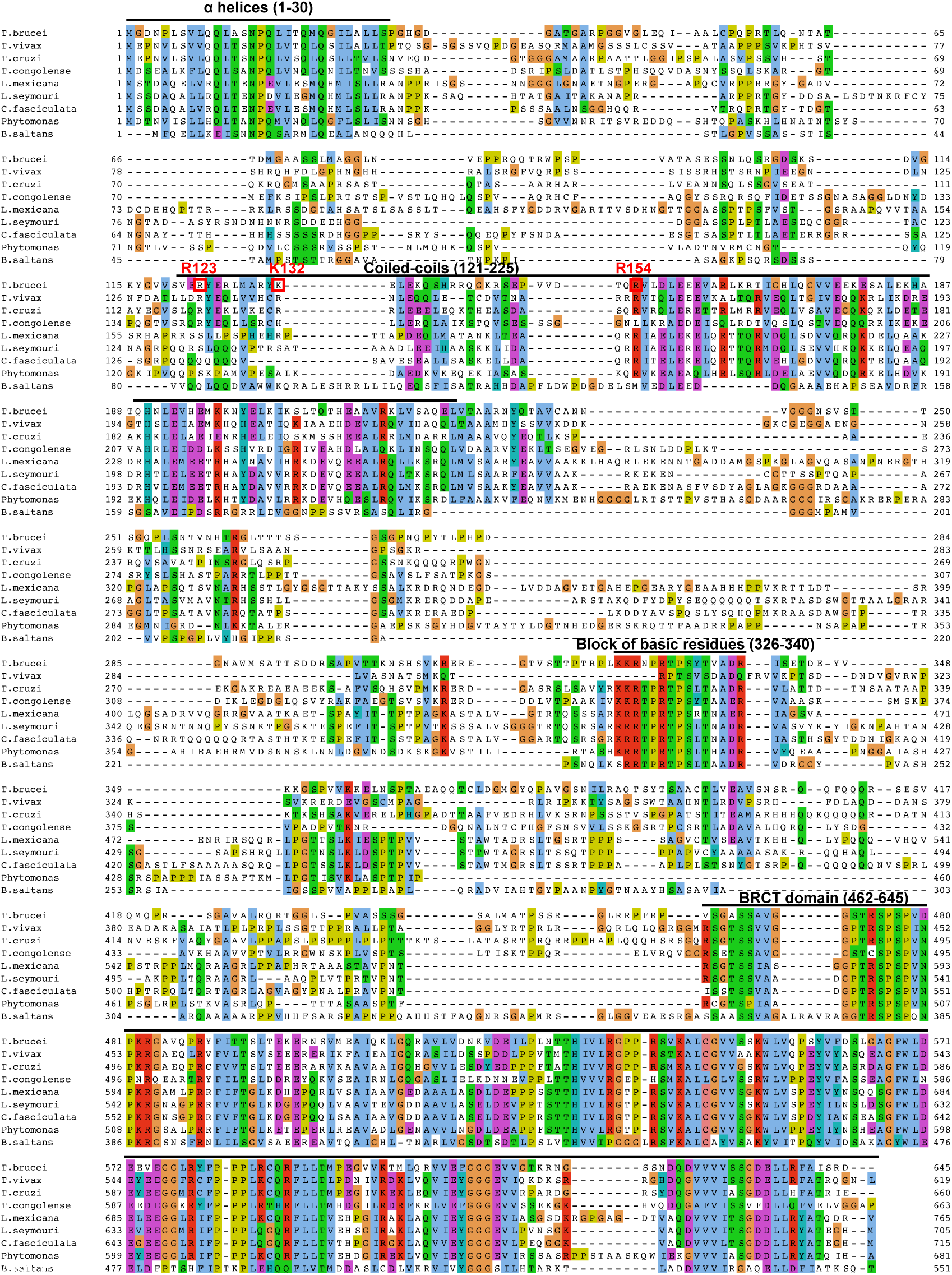
Multiple sequence alignment of KKT4. KKT4 proteins from various kinetoplastids were aligned using MAFFT (L-INS-i) (Katoh and Standley, 2013) and visualized with CLUSTALX coloring scheme in Jalview (Waterhouse et al., 2009). Secondary structure predictions, coiled-coil predictions were performed using PSIPRED (Buchan et al., 2013) and COILS (Lupas et al., 1991), respectively. Charged residues mutated in this study are highlighted in red boxes.

**Figure S3.**
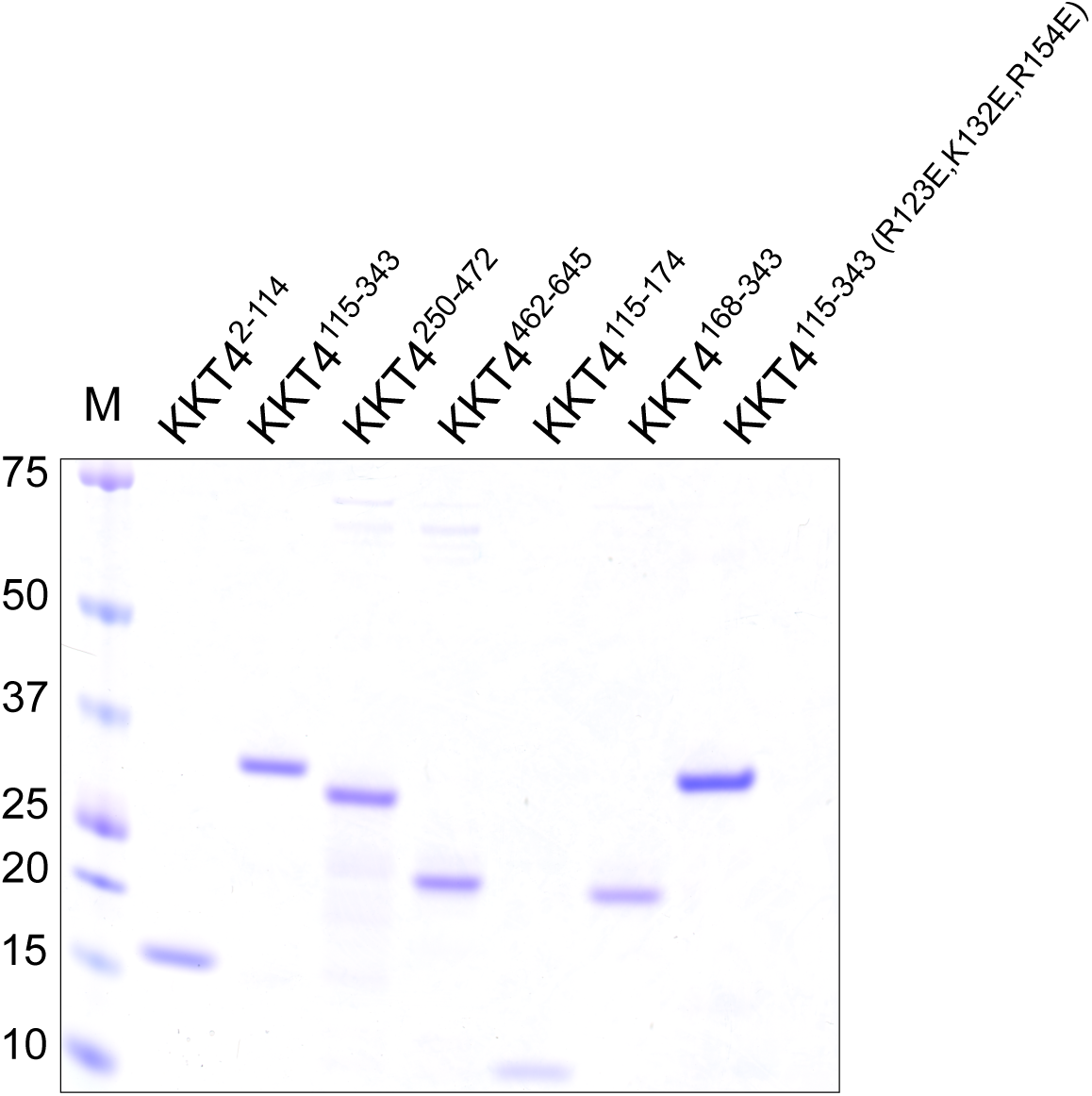
Truncated KKT4 proteins used for microtubule co-sedimentation and laser trap assays. KKT4 fragments fused with an N-terminal 6HIS tag were expressed and purified from *E. coli* and run on SDS-PAGE gel and stained with Coomassie, showing the purity of each sample.

### Supplemental Tables

Table S1.

Summary of optical trap-based bead motility assays, related to Figures 5F and 5G.

**Table S2.** Trypanosome cell lines used in this study.

| Strain | Description | Used in Figure |
| --- | --- | --- |
| SmOxP9 | Parental cell line that expresses TetR and T7 RNAP (Poon et al., 2012) | – |
| BAP665 | TY-YFP-KKT4, TY-tdTomato-KKT2 | 1 |
| BAP943 | TY-YFP-KKT4, TY-tdTomato-MAP103 | 1 |
| BAP1082 | TY-YFP-KKT4, TY-tdTomato-KKT2, KKT4 RNAi | 2 |

**Table S3.**
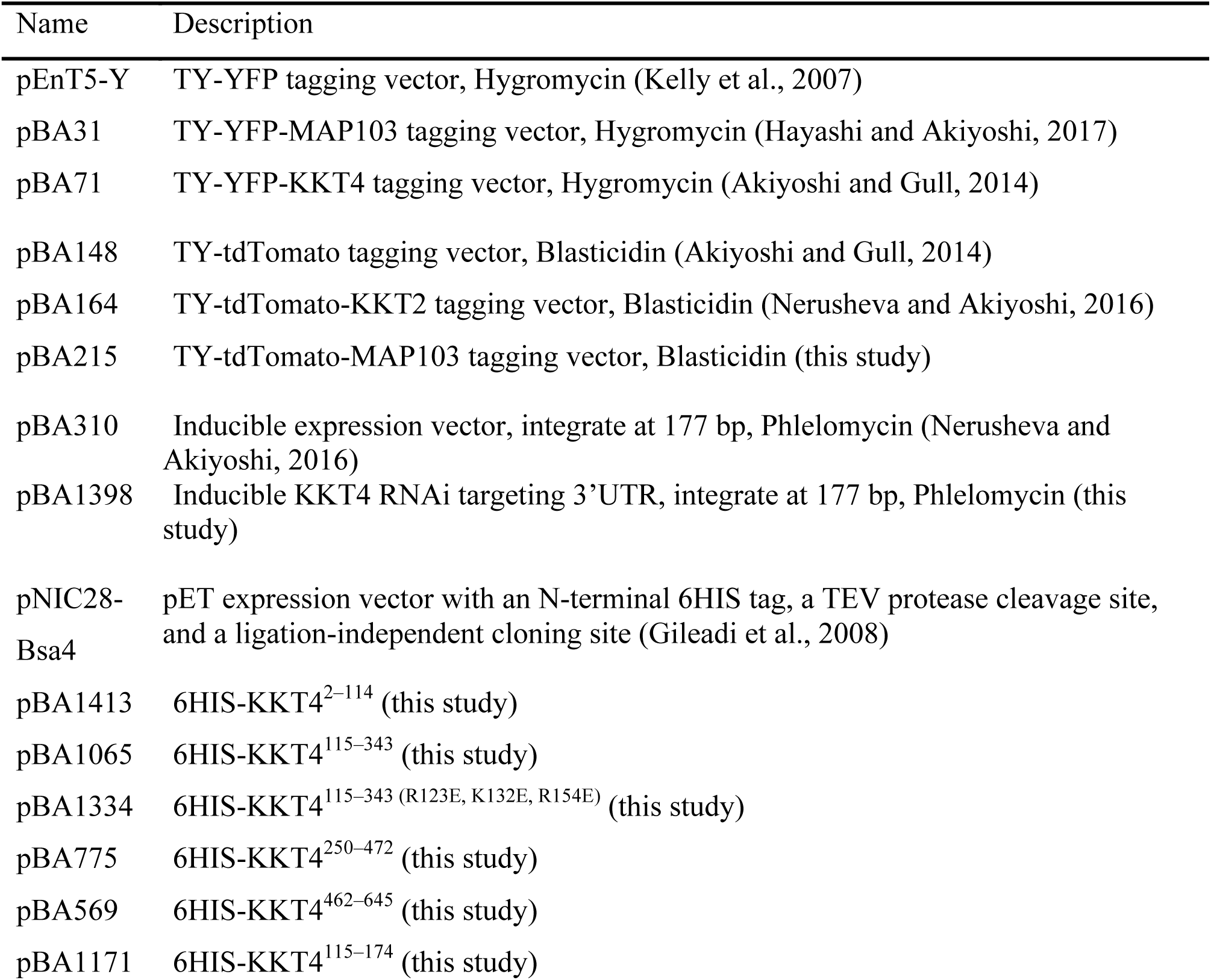
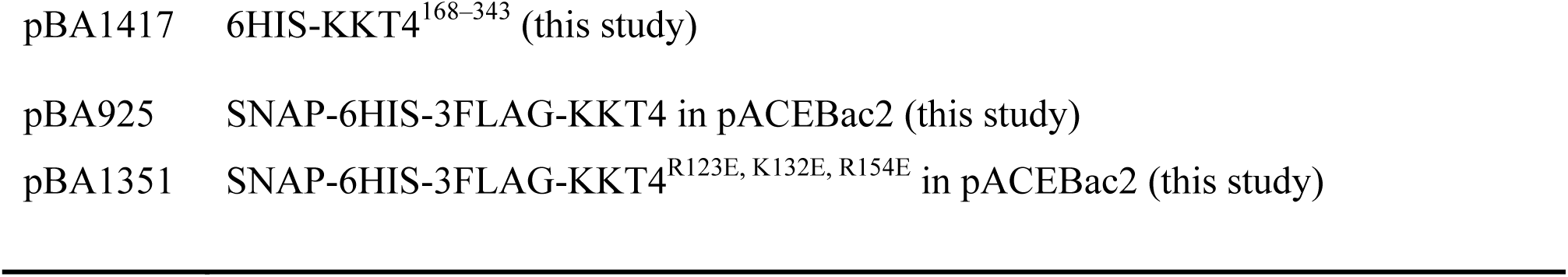
Plasmids used in this study.

**Table S4.**
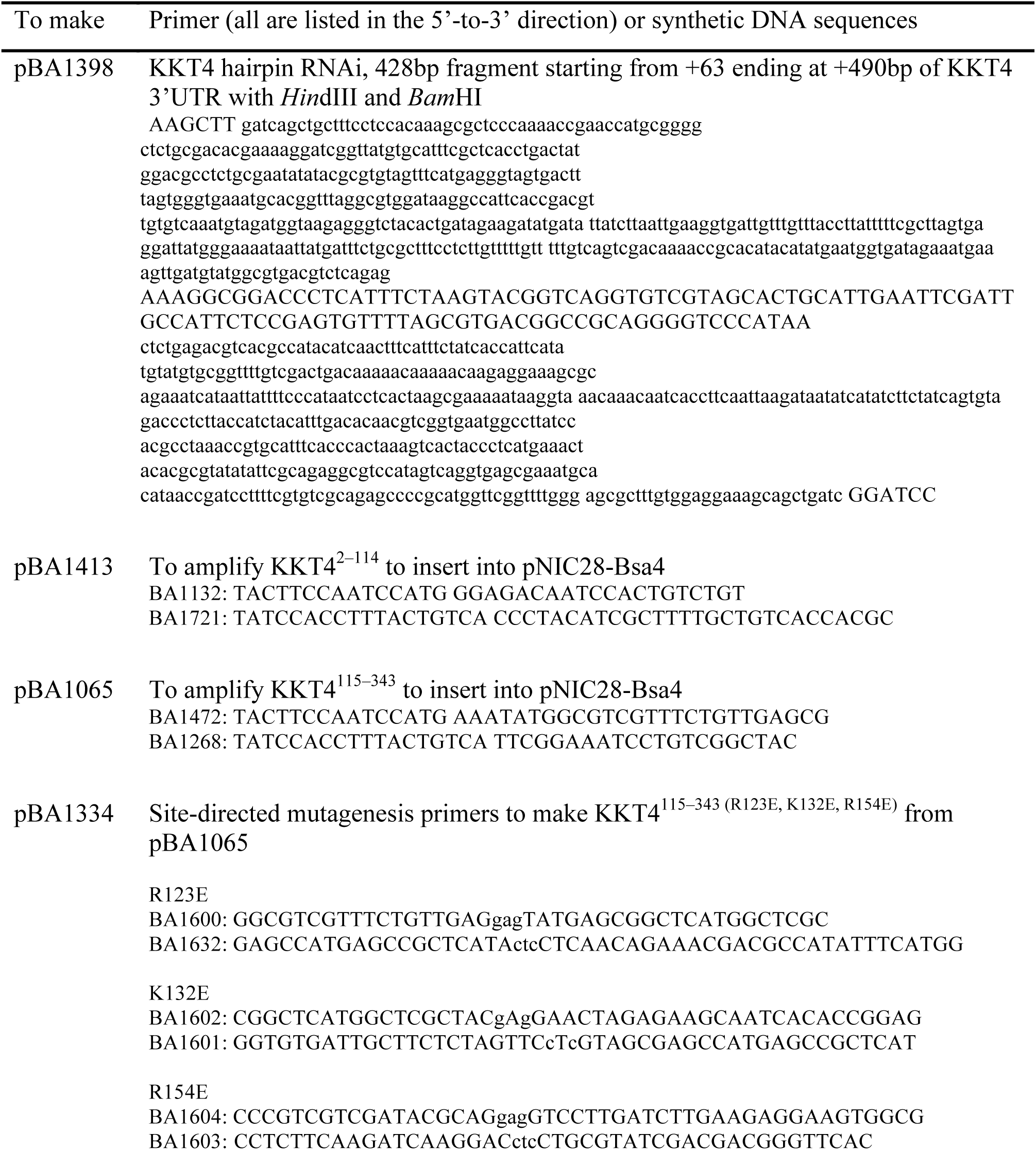
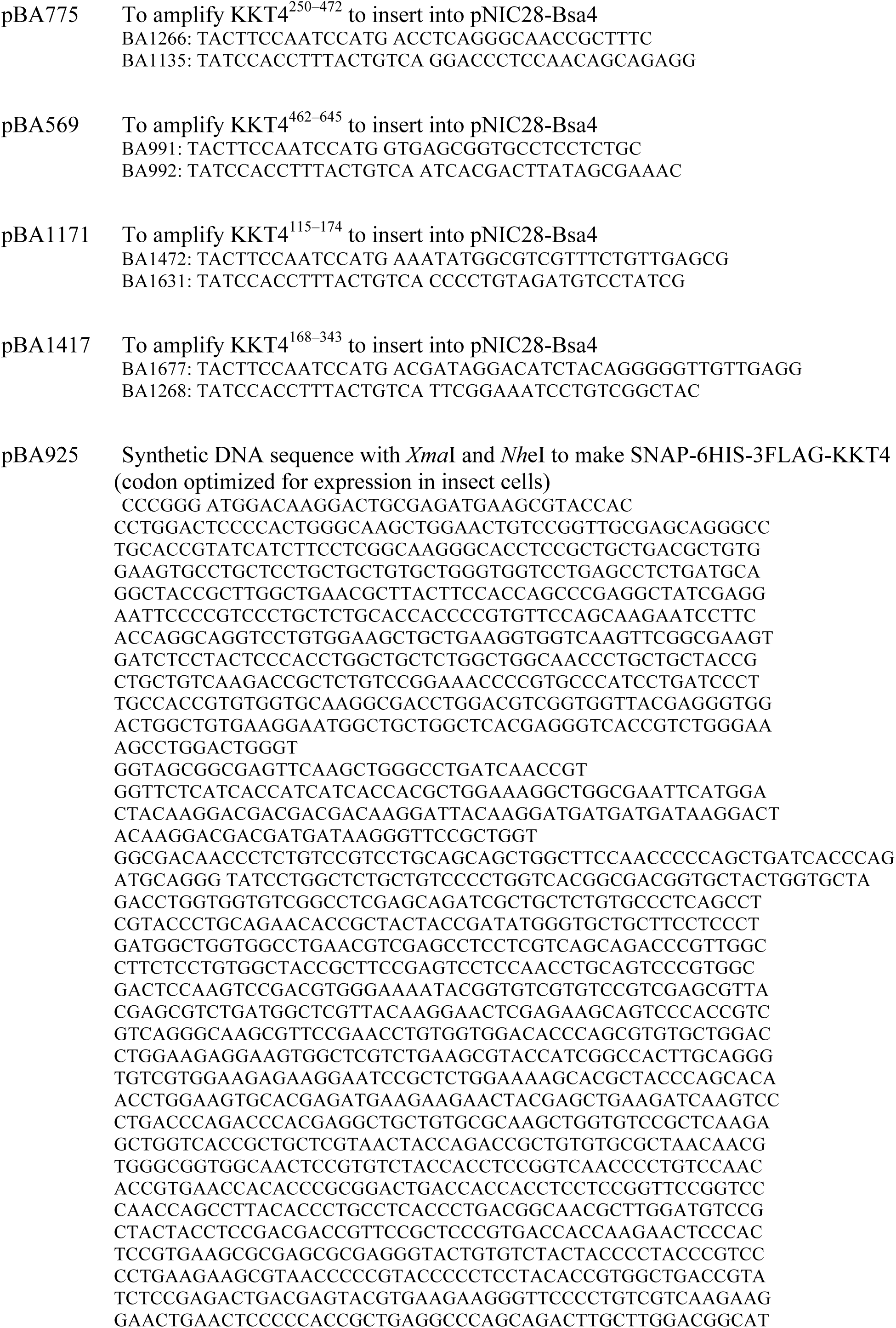
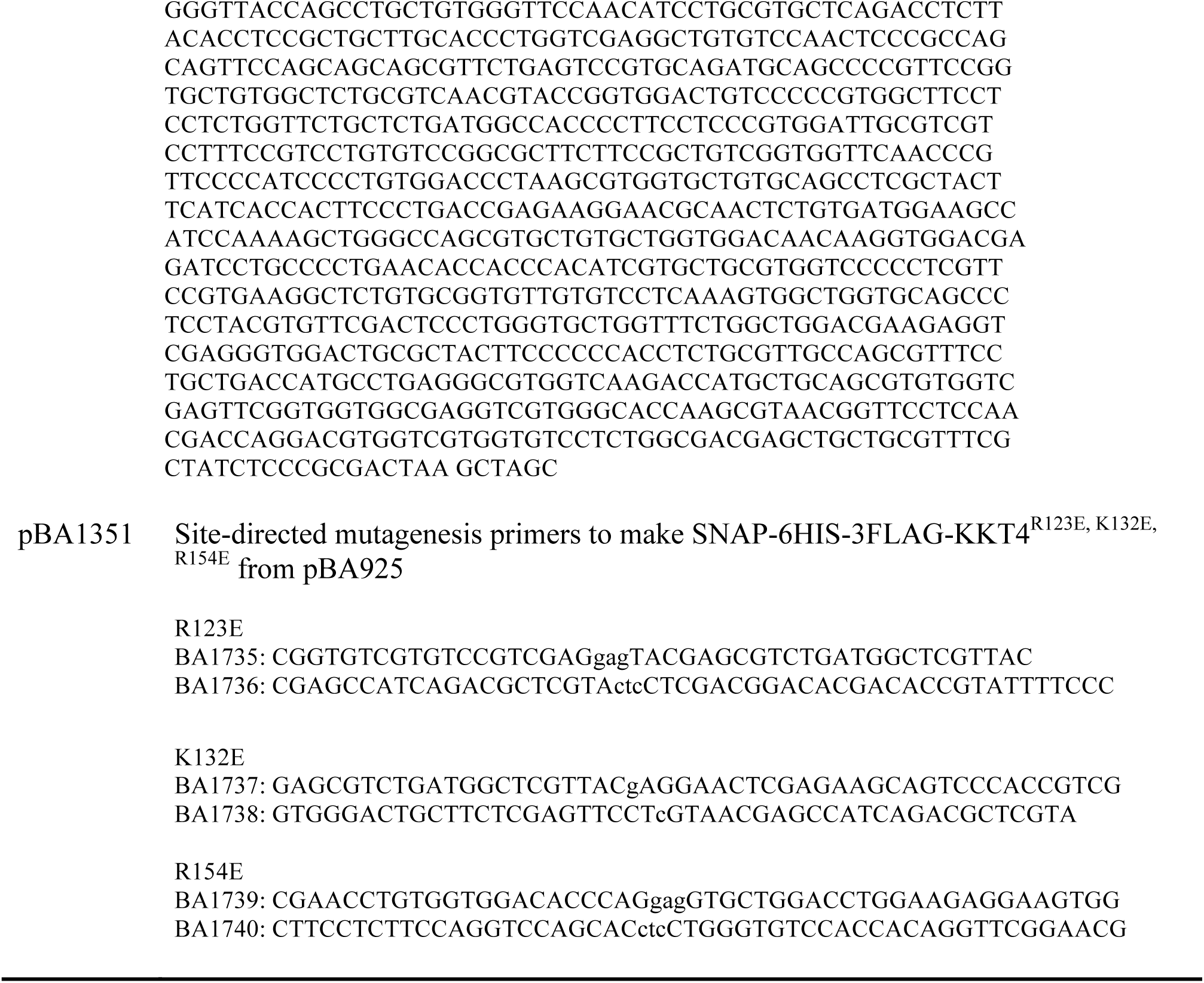
Primers and synthetic DNA sequences used in this study.

